# The strength of sexual signals predicts same-sex paring in termites

**DOI:** 10.1101/2024.03.07.583902

**Authors:** Nobuaki Mizumoto, Sang-Bin Lee, Thomas Chouvenc

**Affiliations:** Department of Entomology & Plant Pathology, Auburn University, Auburn, AL, 36849, USA; Evolutionary Genomics Unit, Okinawa Institute of Science & Technology Graduate University, Onna-son, Okinawa, 904-0495, Japan; Computational Neuroethology Unit, Okinawa Institute of Science & Technology Graduate University, Onna-son, Okinawa, 904-0495, Japan; Entomology and Nematology Department, Ft. Lauderdale Research and Education Center, Institute of Food and Agricultural Sciences, University of Florida, Ft. Lauderdale, FL, 33314, USA

**Keywords:** homosexual behavior, movement coordination, pheromone, same-sex pairing, social insects

## Abstract

Same-sex sexual behavior (SSB) is an enigma in behavioral ecology as it does not result in reproduction. Proximately, the evolution of sexual signals is critical for the evolution of SSB in a sex-specific manner. For signal receivers, the loss of sexual signals leads to smaller phenotypic sex differences, leading to frequent accidental SSB between receivers. Alternatively, for senders, sexual signals could help locate another sender, enhancing intentional SSB as in heterosexual pairing. Here, we demonstrate this link between sex pheromone signaling and the frequency of same-sex pairing in two *Coptotermes* termites that use the same chemical as sex pheromones but in different quantities. In termites, mating pairs engage in tandem runs, where a male follows a female with sex pheromones. We found that female-female tandems were more stable in *C. formosanus,* whose females produce more pheromones, while the male-male interactions were more frequent in *C. gestroi*, whose females produce fewer pheromones. Thus, stronger pheromones lead to sender-sender SSB, while weaker pheromones lead to receiver-receiver SSB. In both species, same-sex tandems were less stable than heterosexual tandems, contrasting with *Reticulitermes*, another termite group that shows frequent same-sex tandems. The proximate mechanism of SSB is diverse, reflecting the heterosexual context.

## Introduction

Same-sex sexual behavior (SSB) is widespread among diverse animals with considerable variations across taxa [1–3]. In most species, the SSB is considered to result from mistaken identity [3,4]. In some cases, SSB provides adaptive value by making the best of a bad job with the shortage of heterosexual partners [5,6]. In either case, the occurrence of SSB is dependent on the mode of mating strategy in heterosexual contexts and strongly affected by the accuracy and carefulness of sex identification of the mating partner [7]. Sexual communication, mediated via sex-specific attracting signals (e.g., sex pheromones), underlies such sexual identification of the other sex. Therefore, even if the SSB does not have adaptive value, the evolutionary patterns of sexual signals could shape the diversity of SSB across species as a by-product. However, the role of sex-specific signals in SSB has remained unexplored.

As signal senders and receivers play distinct roles in mate pairing, the effect of sexual signals on SSB should differ between sender-sender and receiver-receiver pairs. For example, the strength of sexual signals has the opposite effect on the sender-sender SSB and receiver-receiver SSB. In species with weak sex-specific signals, more frequent SSB between receivers is expected compared with those with strong signals [8]. Because it needs more effort for receivers to distinguish senders from receivers with weak sexual signals, receivers may mistake the sex of the partner. On the other hand, more SSB between signal senders could be possible in the species with strong signals. It is rarer to observe SSB between senders as they are usually the passive sex during paring [9]. Nevertheless, if the signals are strong enough, it should be easier for even senders to find other senders than other receivers. Thus, the strength of sexual signals is expected to modify the relative occurrence of receiver-receiver and sender-sender SSB across species.
Mate pairing in termites provides an ideal model system to study the evolution of SSB. Termites form life-long monogamous pairs to establish colonies [10]. During a brief period, alates (winged adults) disperse from their nests. Both females and males land on the ground or trees, shed their wings, and run to search for a mating partner. Upon joining, a pair performs a tandem run. In neoisopteran termites, the male always follows the female that produces sex pheromones [11], maintaining contact in a highly coordinated manner while seeking a suitable site for colony foundation. Although tandem running involves communication via sex pheromones, same-sex tandem running can be observed in either sex [11,12]. Sex pheromones should play different roles between female-female pairs and male-male pairs. In male-male pairs, same-sex tandem can happen once one male starts following another male. Thus, in the case of male-male tandems, SSB can happen because of mistaken identity. This implies that species with weak sex pheromones may provoke male-male tandem pairing. On the other hand, female-female tandem should not come from the mistaken identity because the sex role is fixed (females do not follow males) [11,13], and one female needs to change the sex role before the initiation of female-female tandem runs when not approached by males. This implies that female-female tandem runs could be intentional. Thus, conversely, the strong pheromone signal of females instead facilitates same-sex tandems, where females can easily follow another female.

In this study, we compared the same-sex tandem running behavior in two *Coptotermes* termites: *Coptotermes formosanus* Shiraki and *Coptotermes gestroi* (Wasmann). These two species share the same chemical for sex-pairing pheromones ((3Z,6Z,8E)-dodeca-3,6,8-trien-1-ol) produced by female tergal glands [14] and were previously observed forming hetero-specific tandem runs [15,16]. However, the quantity of pheromones differs between these two species, where *C. formosanus* females produce ten times more pheromones than *C. gestroi* females [14]. Due to this signal difference, we here expect that male-male tandem is more frequent in *C. gestroi* than *C. formosanus*, while female-female tandem is more frequent in *C. formosanus* than *C. gestroi*. Finally, when a pair is separated during a heterosexual tandem run, both species display sexual dimorphism, with females pausing to wait for males and males moving around to search for females. [17,18]. We, therefore, also investigated if an individual could display the typical behavior of the opposite sex in interrupted same-sex tandem runs, as shown in another termite lineage [11].

## Methods

### Termites and Experimental Arena

We collected alates using a light-trapping system [19] at dusk between 27-29 March for *C. gestroi* and 21-22 April, 1-2 May for *C. formosanus* in 2021 in Broward County (Florida, USA) during synchronized dispersal flights. We collected all the alates at a single site. We brought the alates to the laboratory and maintained them on wet cardboard at 28°C. We used individuals who did not shed their wings by themselves to prevent any prior experience with tandem runs. After inducing shedding wings, we observed their behaviors within 12 h. We used each termite only once.

We performed all observations in an experimental arena made of a Petri dish (ø = 150 mm) filled with moistened plaster. The Petri dish had a clear lid during observations. A video camera (HC-V770, Panasonic, Osaka, Japan) was above the arena, adjusted so that the arena filled the camera frame. We introduced a pair of termites into the arena. Each pair was recorded for 30 minutes in 30 frames per second (FPS). We obtained 49, 61, and 44 videos for female-male, female-female, and male-male in *C. formosanus*, and 40, 40, and 45 videos in *C. gestroi*. We extracted the coordinates of the centroids of termite movements from all obtained videos using the video-tracking system UMATracker [20]. We down-sampled all coordinates to a rate of five FPS for subsequent analyses. We measured the diameter of the dish and body length of two termites in pixels for each video using a Python program.

### Tandem analysis

To compare the duration of tandem runs between pair combinations and species, we automatically determined that pairs were in interactions when the distance between their centroids was within the sum of their body length. We did not count short interactions (< 5 seconds) as interactions and short separations (< 2 seconds) during interactions as interruptions. Then, for each interaction event, we estimated the leader by checking the relative directions of partners and their movement directions. We calculated the dot products of these two vectors and defined whether each partner was leading the other (dot vector < 1, the partner exists behind the moving direction) or not (dot vector ≧ 1). During each interaction event, we regarded termites in tandem only when one individual was the predominant leader, where the proportion of time being a leader for one individual was more than 50% of the total interaction duration. This treatment excluded non-tandem interactions, such as inspecting the partner by antennation or staying together while inspecting a specific area of the arena.

We compared the proportion of the total time spent in interactions, tandem runs, and non-tandem interactions across different pairing combinations (heterosexual, female–female, or male–male) for each species, or between species for each pairing combination. We transformed proportional data using logit transformation after adding 0.01 to the observed proportions to avoid infinite values [21]. We used the Welch t-tests to compare species for each pair combination, while one-way ANOVA with Tukey’s HSD to compare combinations for each species. In the t-test and Tukey’s HSD, we obtained Cohen’s d value as effect sizes [cohens_d() function in the rstatix package in R]. We also studied how tandem runs were stable once they started by comparing the duration of each tandem running event. We used mixed-effects Cox models [coxme() function in the coxme package in R [22]], with pair combination (or species) treated as a fixed effect and each pair id as a random effect. We compared the durations between species for each combination and among combinations for each species separately.

Finally, for each separation event after tandem running events, we obtained the movement speed of both leader and follower termites after 5 seconds of separation to compare the movement speed between them, using a paired t-test with Cohen’s d value as effect sizes. We also calculated the absolute difference in movement speed. We compared the difference in movement speed using linear mixed-effects models [lmer() function in the lme4 package in R [23]], using the same approach as the above mixed-effect Cox models.
All data analyses were performed using R v4.3.0 [24], and source codes are available on GitHub (https://github.com/nobuaki-mzmt/cop_homo_tandem_cf-vs-cg).

## Results

The species difference in tandem running behavior depended on the pairing combinations. In heterosexual pairing, *C. formosanus* spent longer time for tandem runs than *C. gestroi* (t-test, *t*_88.0_ = 4.30, *P* < 0.001, *d* = 0.90, Figure 1C) or interactions (t-test, *t*_84.3_ = 5.38, *P* < 0.001, *d* = 1.14, Figure 1B), but not for non-tandem interactions (t-test, *t*_88.0_ = 1.04, *P* = 0.299, *d* = 0.22, Figure 1D). Once they started tandem runs, *C. formosanus* showed more stable tandems (mixed-effects Cox model, 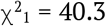, *P* < 0.001, Figure S1). Note that this result contradicts a previous study that detected no significant difference between these two species [16], and this contradiction is discussed in the supplementary material (see Supplementary Text S1, Figure S2, and Table S1). A similar pattern was observed in female-female pairing, where *C. formosanus* spent longer time for tandem runs (t-test, *t*_96.4_ = 8.62, *P* < 0.001, *d* = 1.65, Figure 1C) and showed more stable tandems than *C. gestroi* (mixed-effects Cox model, 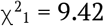, *P* = 0.002, Figure S1). Note that time spent in interactions was not different (t-test, *t*_83.6_ = 1.62, *P* = 0.11, *d* = 0.33, Figure 1B) because *C. gestroi* expended more time for non-tandem interactions (t-test, *t*_75.2_ = 5.67, *P* < 0.001, *d* = 1.17, Figure 1D). On the other hand, male-male pairing showed a distinct pattern, where *C. gestroi* spent longer for same-sex interactions than *C. formosanus* (t-test, *t*_73.9_ = 6.36, *P* < 0.001, *d* = 1.35, Figure 1B). The most social interactions in *C. gestroi* did not end up tandem runs as there was no difference in the time spent in tandem runs between species (t-test, *t*_78.5_ = 0.93, *P* < 0.353, *d* = 0.20, Figure 1C), but *C. gestroi* expended much more time for non-tandem interactions (t-test, *t*_84.9_ = 7.43, *P* < 0.001, *d* = 1.58, Figure 1D). Interestingly, once tandem started, *C. formosanus* showed more stable tandems than *C. gestroi* (mixed-effects Cox model, 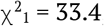, *P* < 0.001, Figure S1). However, in either species, same-sex tandem runs were much less stable than heterosexual pairing, with larger difference between female-female pairs and male-male pairs in *C. formosanus* (mixed-effects Cox model, 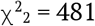, *P* < 0.001; Tukey’s HSD, FM-FF: *P* < 0.001; FM-MM: *P* < 0.001; FF-MM: *P* < 0.001; Figure S1), while no difference between female-female pairs and male-male pairs in *C. gestroi* (mixed-effects Cox model, 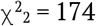, *P* < 0.001; Tukey’s HSD, FM-FF: *P* < 0.001; FM-MM: *P* < 0.001; FF-MM: *P* = 0.839; Figure S1). The same pattern was observed in the time spent in tandem runs (Figure 1C).

**Figure 1.**
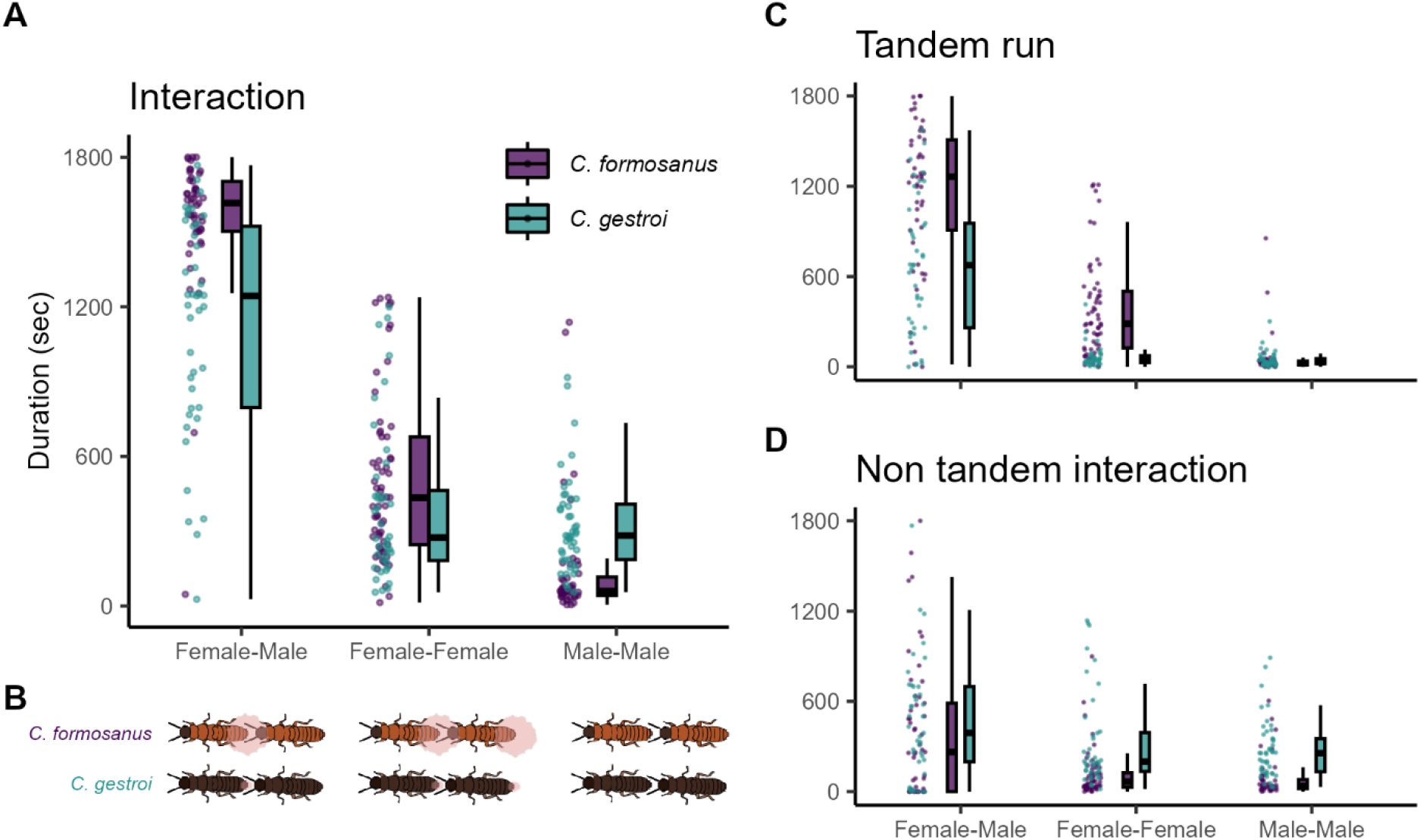
Comparison of the total time spent in mating interactions among pairing combinations and species. The proportion of time spent in interactions (A) was further divided into tandem running (C) and non-tandem interactions (D). (B) The sex-attracting pheromone is visualized. The data is bound to 1,800 sec as the observation ended at this point.

After separations, combinations with stable tandem runs showed clear speed differences between partners (Figure 2). As shown in previous studies [17,18], in heterosexual pairs, male followers sped up while female leaders slowed down after separation to enhance the probability of reunions in both *C. formosanus* (t-test, *t*_366_ = 16.9, *P* < 0.001, *d* = 1.60) and *C. gestroi* (t-test, *t*_438_ = 15.7, *P* < 0.001, *d* = 1.34). Similar movement differences between partners were observed in the female-female pairing of *C. formosanus* (t-test, *t*_1208_ = 20.1, *P* < 0.001, *d* = 1.15), which shows stable tandem runs (Figure 1C, S1). However, in other same-sex pairs, the speed difference between leaders and followers after separation was small and had negligible effect sizes; male-male pair in *C. formosanus* (t-test, *t*_278_ = 0.03, *P* = 0.978, *d* = 0.003), female-female pair in *C. gestroi* (t-test, *t*_300_ = 3.62, *P* < 0.001, *d* = 0.39), and male-male pair in *C. gestroi* (t-test, *t*_353_ = 2.13, *P* = 0.034, *d* = 0.22). Similarly, the levels of difference between partners were smaller in same-sex pairing, compared with heterosexual pairs both in *C. formosanus* (LMM, 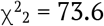, *P* < 0.001; Tukey’s HSD, FM-FF: *z* = 4.26, *P* < 0.001; FM-FF: *z* = 8.57, *P* < 0.001; FM-FF: *z* = 5.69, *P* < 0.001) and *C. gestroi* (LMM, 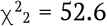, *P* < 0.001; Tukey’s HSD, FM-FF: *z* = 5.65, *P* < 0.001; FM-FF: *z* = 6.56, *P* < 0.001; FM-FF: *z* = 0.844, *P* = 0.675).

**Figure 2.**
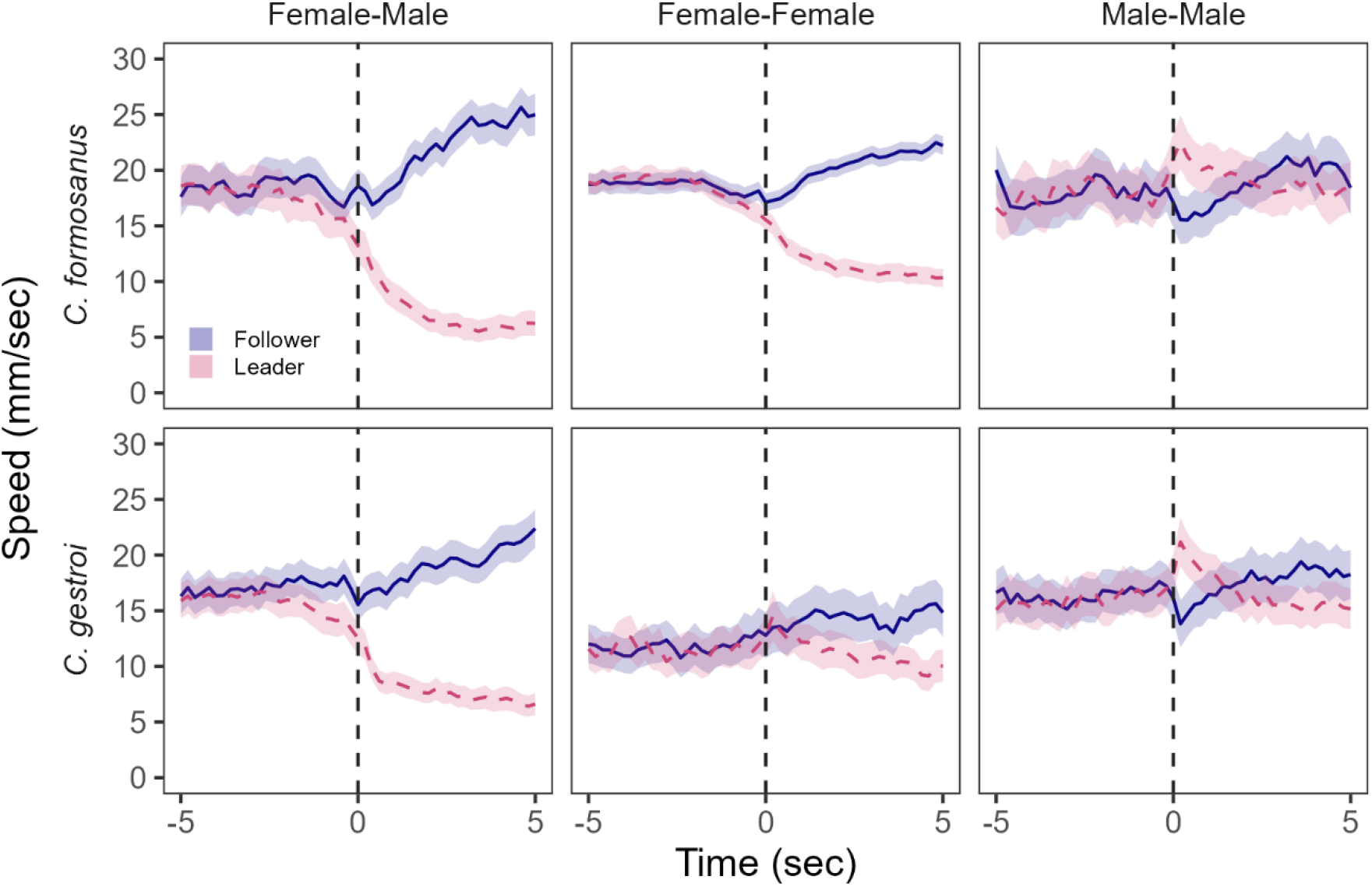
Movement speed of termites before and after separation from tandem running. Separation timing (time = 0 sec) was determined when the distances between partners were larger than the sum of their body lengths. Solid lines indicate followers, while dashed lines indicate leaders. Lines and shaded regions indicate mean ± 95% confidence intervals.

## Discussion

We found an association between same-sex pairing behavior and sex pheromone quantity in two termite species with shared chemicals. First, female-female pairing was more stable in *C. formosanus*, the species with more sex pheromones. Because females are leaders in heterosexual pairs of *Coptotermes* termites, females need to decide to play follower roles before the initiation of same-sex pairing [11], where females could use sex pheromones to maintain stable movement coordination with same-sex individuals. On the other hand, although both species showed little male-male tandem running, *C. gestroi* males, the species with a smaller quantity of sex pheromones, spent more time interacting with same-sex partners. Male-male pairings of these species could be driven by mistaken identity, where *C. gestroi* required more time to inspect the partner’s sex due to smaller sexual dimorphism. Either way, our study clearly demonstrated that the evolution of SSB in termites is inseparable from the evolution of sex-attracting signals.

By comparing with other termite lineages, our results suggest that the same-sex tandems in *Coptotermes* termites are not functional behaviors but could exist because they inherited them from the ancestral state of termite tandem running behavior [11], as in the framework of the previous work [4]. Previous studies on same-sex tandem runs in termites have focused only on *Reticulitermes* termites [11,12,25,26], which belong to the same family as *Coptotermes*. In *Reticulitermes* termites, same-sex pairing functions by providing survival benefits [5,12,27] and is not a result of mistaken identity [11]. However, a distinct pattern observed in *Coptotermes* termites suggests that same-sex tandem in *Coptotermes* termites is rather a behavior expressed outside of its original function. In *Reticulitermes* termites, same-sex tandem pairs were as stable as heterosexual tandem pairs, and one partner behaves like the other sex upon separation [11]. In *Coptotermes* termites, on the other hand, even once they form a same-sex tandem pair, both female-female and male-male tandems were far less stable than heterosexual tandems (Figure 1, S1). Furthermore, upon separation, movement dimorphisms between partners, a key for efficient reunion, were not as strong as heterosexual pairs or even absent (Figure 2). These all indicate that *Coptotermes* termites do not tend to adjust their movement patterns to maintain same-sex tandems actively. The exception is *C. formosanus* female-female pairing, which may have an unknown behavioral function.

Even though the same-sex pairing of *Coptotermes* termites is not adaptive, our study on their interspecific variations shows the mechanical aspects of the evolution of SSB. For example, a previous theoretical study predicted that SSB is caused by the absence of perfect sex discrimination, where the loss of sexual signals and indiscriminate mating coevolve together [28]. Our result supports this idea by showing that the species with weaker sexual signals showed more frequent same-sex interactions than the other. In other words, *C. gestroi* has a broader mating filter than *C. formosanus* and tries to pair with non-female individuals, as shown in a cricket [29], a burying beetle [30], and a water strider [31]. Furthermore, although all of the previous studies have considered the SSB between individuals of signal-receiving sex (typically males) [4,7,28], our results also indicated that the evolution of sexual signals could also affect the evolution of SSB between signal-sending sex (typically females), where stronger signals might attract same-sex individuals and lead to sender-sender SSB. This newly suggested relationship needs further empirical tests in other animal lineages, possibly in the context of variable densities of available partners.

In summary, our study highlights the diversity of SSB that can exist even within a closely related species. By connecting mating behavior in heterosexual contexts with the occurrence of SSB across species, comparative behavioral analysis has the potential to answer the questions relating to the evolution of SSB.

## Data accessibility

All data and source codes for analyzing them are available at Github: https://github.com/nobuaki-mzmt/cop_homo_tandem_cf-vs-cg., and the accepted version will be deposited at Zenodo.

## Authors’ contributions

NM: Conceptualization, Methodology, Formal analysis, Data curation, Writing – original draft

SBL: Methodology, Investigation, Data curation, Writing – review & editing

TC: Resources, Writing – review & editing

## Acknowledgments

We thank Aoi Mizumoto for her assistance during the video analysis. This work was supported by a JSPS Research Fellowship for Young Scientists CPD to NM (20J00660), a Grant-in-Aid for Early-Career Scientists (21K15168) to NM, and an IPSF fellowship from OIST to NM.

