## Supplementary material for "The strength of sexual signals predicts same-sex paring in termites": Text S1, Figure S1-2, Table S1

This file includes:
Text S1
Figure S1-2
Table S1

### Supporting Information Text

**Text S1. Comparison of heterosexual tandems across experimental years.**
This study detected a clear difference in stability and time spent on heterosexual tandem running behavior between *C. formosanus* and *C. gestroi* (Figure 1, S1). However, this result contradicts the previous study that found a similar level of tandem stability between these two species (see Figure 1 of [16]). To investigate the source of this inconsistency, we reanalyzed the data of previous studies using *C. formosanus* and *C. gestroi* in this region [16,18] using the same methodology as this study. We have three different datasets: one obtained in 2019 only for *C. gestroi* to investigate their density-dependent behavioral change, one obtained in 2020 for both species to investigate the heterospecific tandem runs, and one obtained in 2021 for both species in this study (Table S1).

In *C. formosanus*, there was no difference between this study and a previous study in the duration of each tandem run (mixed-effects Cox model, $\chi^2_1 = 0.132$, $P = 0.716$; Figure S2A) and total time spent in tandem runs (t-test, $t_{15.7} = 0.162$, $P = 0.873$, $d = 0.052$; Figure S2B). On the other hand, in *C. gestroi*, we found different stabilities of tandem runs across studies, where results obtained in 2020 showed marginally higher duration of tandem runs compared to others (mixed-effects Cox model, $\chi^2_2 = 6.36$, $P = 0.042$; Tukey's HSD, 2020-2019: $z = 1.90$, $P = 0.134$, 2021-2019: $z = 0.815$, $P = 0.688$, 2021-2020: $z = 2.52$, $P = 0.031$; Figure S2A). Note that the total time spent in tandem runs was not different among studies (ANOVA, $F_2 = 0.51$, $P = 0.603$; 2020-2019: $d = 0.316$, 2021-2019: $d = 0.013$, 2021-2020: $d = 0.417$; Figure S2B). When we compared the tandem running behavior between two *Coptotermes* termites by accounting for the experimental years as a random effect, *C. formosanus* showed higher stability (mixed-effects Cox model, $\chi^2_1 = 45.6$, $P < 0.001$, Figure S2A) and a longer period of tandem runs than *C. gestroi* (linear mixed-effects model, $\chi^2_1 = 20.8$, $P < 0.001$, Figure S2B). Thus, tandem running stability is actually different between *C. gestroi* and *C. formosanus*, but Mizumoto et al., 2021 [16] failed to detect the difference probability due to the smaller sample size (Table S1).

In *C. gestroi*, why did the experiment in 2020 show higher stability of tandem runs compared to 2019 and 2021, even if the difference is small? This experiment focused on the interspecific tandem runs between *C. gestroi* and *C. formosanus*; thus, they observed their tandem runs on the date when both species swarmed together, which needs to be the very end of the swarming season of *C. gestroi* and the beginning of that of *C. formosanus* [15] (Table S1). As physiological conditions of termite alates can change across the swarming seasons in both *C. gestroi* and *C. formosanus* [32], this variability of the swarming season might have changed their tandem running behavior, too. Further studies need to clarify this hypothesis.

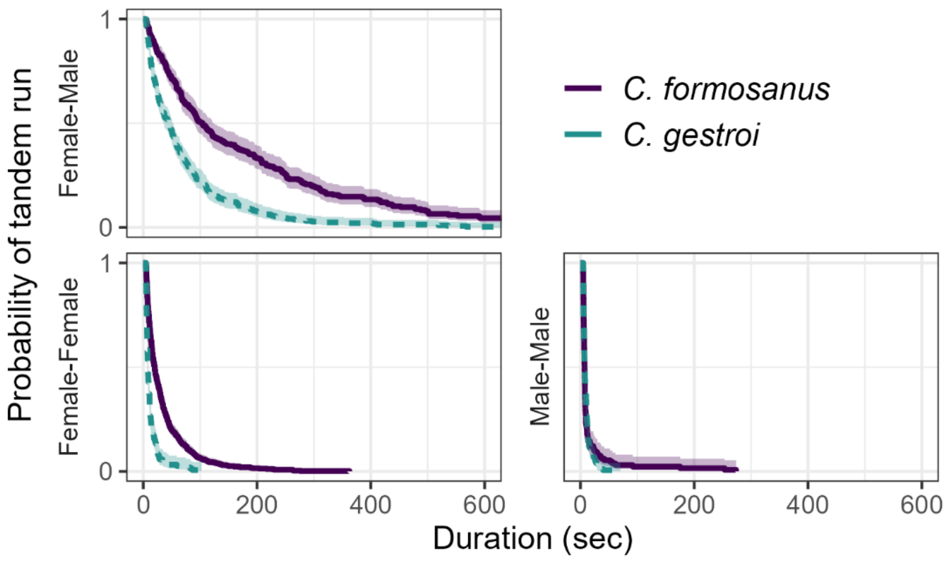


**Figure S1.** Comparison of the duration of tandem running until separation across different pairing combinations and species. Kaplan–Meier survival curves were generated for each pairing combination. The marks for censored data are not shown. Shaded regions show 95% confidence intervals.

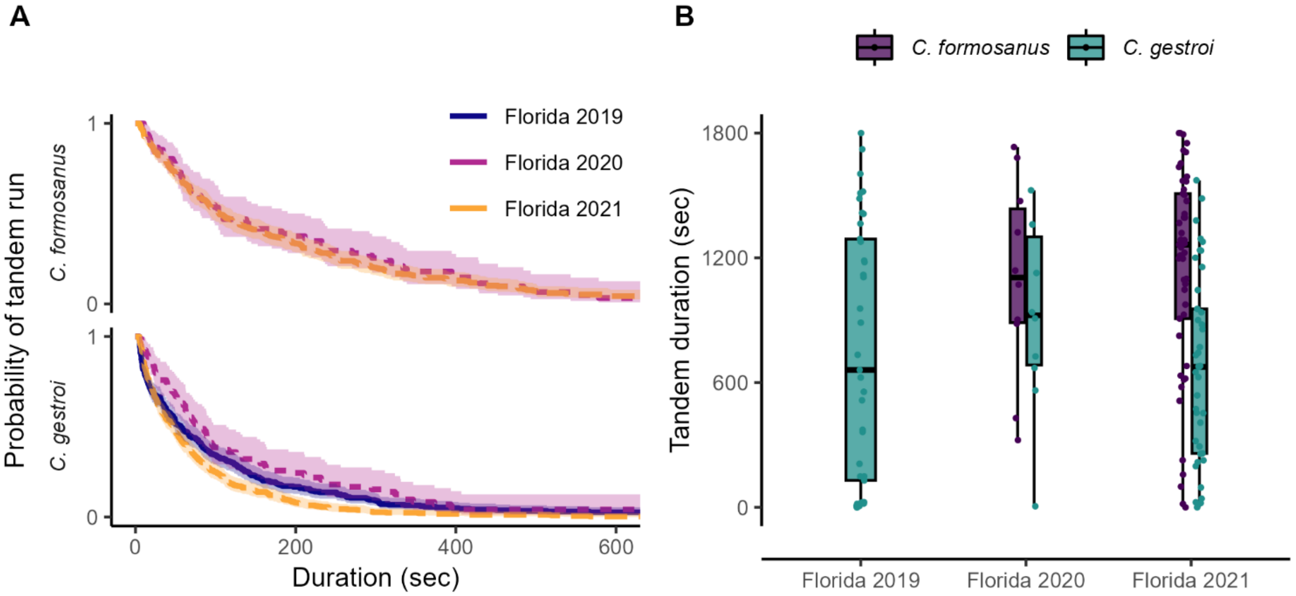


**Figure S2.** Comparison of tandem running behavior in heterosexual pairs across different experiments in (A) the duration of tandem running until separation and in (B) total time spent in tandem during 30-minute observation. Data from Florida 2019 is from Mizumoto et al., 2020 [18], from Florida 2020 is from Mizumoto et al., 2021 [16], and Florida 2021 is from this study. (A) Kaplan–Meier survival curves were generated for each pairing combination. The marks for censored data are not shown. Shaded regions show 95% confidence intervals.

**Table S1**. Comparison of experimental conditions across studies.

| Species | Experiments | Date | Tandems | Pairs | Temp | Arena | Ref |
|---|---|---|---|---|---|---|---|
| *C. gestroi* | Florida 2019 | March 5,8 | 484 | 37 | 28 | 140 | [18] |
| *C. gestroi* | Florida 2020 | April 18,20 | 89 | 10 | 28 | 140 | [16] |
| *C. gestroi* | Florida 2021 | March 27-29 | 721 | 40 | 28 | 150 | This study |
| *C. formosanus* | Florida 2020 | April 18,20 | 94 | 10 | 28 | 140 | [16] |
| *C. formosanus* | Florida 2021 | April 21-22, May 1-2 | 421 | 49 | 28 | 150 | This study |

Tandems: the number of tandem running events used for survival analysis (Figure S2). Arena indicates the size of arena in mm.
